# SAMA: a contig assembler with correctness guarantee

**DOI:** 10.1101/2024.07.10.602853

**Authors:** Leena Salmela

## Abstract

In genome assembly the task is to reconstruct a genome based on sequencing reads. Current practical methods are based on heuristics which are hard to analyse and thus such analysis is not readily available. We present a model for estimating the probability of misassembly at each position of a de Bruijn graph based assembly. Unlike previous work, our model also takes into account missing data. We apply our model to produce contigs with correctness guarantee. Our model may have further applications in downstream analysis of contigs or in any analysis working directly on the de Bruijn graph. Our experiments show that when the coverage of *k*-mers is high enough, our method produces contigs with similar contiguity characteristics as state-of-the-art assemblers which are based on heuristic correction of the de Bruijn graph.

## 1 Introduction

Genome assembly is a classical problem in computational biology where the task is to reconstruct a genome based on sequencing reads. Theoretically, genome assembly has been formulated as finding the shortest common superstring of the reads, finding a Hamiltonian path in a string graph, or finding a Eulerian tour in a de Bruijn graph. However, the solutions to these problems are almost never unique, and thus practical methods are instead based on heuristics. These heuristics are hard to analyse and thus such analysis is not readily available [4]. Furthermore, it has been shown that misassemblies are present in the output sequences due to not considering structural correctness properly [10].

Current solutions produce a set of sequences that are claimed to be substrings of the genome. Most assemblers are based on unitigs which are strings corresponding to nonbranching paths in a string graph or a de Bruijn graph and thus guaranteed to be substrings of the genome if there is no missing data, i.e. missing branches in the graph. A further theoretical development is to produce omnitigs [14] which are guaranteed to be substrings of each genome that could produce the string graph or de Bruijn graph but again it is assumed there is no missing data. In the presence of missing data, these approaches may report strings that are not substrings of the original genome.

Several works have previously investigated under which conditions a genome assembly is correct. It has for example been shown that if enough information is available, a unique and correct solution can be found in polynomial time [8, 12]. Enough information can be characterized as having enough information for the solution to be unique [12] or requiring that all repeat instances of the genome are spanned by some read [8]. However, the requirement for enough information essentially means that there can be no missing data and these approaches do not easily extend to estimating the probability of correctness for a certain position in the assembly.

Probabilistic frameworks have previously been applied to genome assembly to produce longer contigs. Medvedev and Brudno [5] and Myers [7] have proposed to report the sequence with maximum likelihood given the reads as the assembled genome. However, this approach does not assign probabilities to misassemblies nor does it consider missing data. Finally, the correctness of genome assembly has been studied in the context of validating the produced assembly [11].

We present a model for structural correctness in de Bruijn graph based assembly. Our model estimates the probability of misassembly for each edge in the de Bruijn graph. We apply our model to de Bruijn graph based assembly to create an assembled genome with a correctness guarantee and a correctness estimate for each position in the assembly. We call the resulting assembler Sequence Assembly avoiding MisAssemblies (SAMA). Although here we apply the model to the genome assembly problem, the model can be of independent interest beyond assembly and could be useful for any downstream analysis based on the results of assembly or to any analysis performed directly on a de Bruijn graph. Our experiments show that when *k*-mer coverage is high enough for computing accurate estimates, our method produces as contiguous assemblies as a state-of-the-art assembler based on heuristic correction of the de Bruijn graph such as tip and bulge removal. Our method is also competitive in runtime and memory usage.

SAMA is available at https://github.com/LeenaSalmela/SAMA/.

## 2 Methods

### 2.1 Definitions and problem formulation

Let us consider a set of reads *R* from a genome *G*. We denote by *r* the number of reads and by *N* the length of the genome. We assume that the reads are sampled at random from the genome. We denote the abundance or frequency of a sequence *S* in the read set *R* by *f* (*S*). When we want to emphasize that the abundance is a constant we use *a* to denote the abundance of a sequence.

A *k*-mer is a sequence of length *k*. The *k*-mer spectrum of a read set is the set of all *k*-mers occurring in the read set. We construct the de Bruijn graph for a read set by enumerating its *k*-mer spectrum. Each *k*-mer occurring in the read set is a node of the de Bruijn graph. There is an edge between two *k*-mers *S*[1 … *k*] and *T* [1 … *k*] if they overlap by *k* − 1 characters, i.e. *S*[2 … *k*] = *T* [1 … *k* − 1] and the *k* + 1-mer *S*[1 … *k*]*T* [*k*] = *S*[1]*T* [1 … *k*] also occurs in the read set. An assembly in a de Bruijn graph consists of a set of paths. Each path *S*_1_*S*_2_ … *S*_*n*_ corresponds to a sequence consisting of the first *k*-mer in the path concatenated with the last characters of each of the remaining *k*-mers: *S*_1_[1 … *k*]*S*_2_[*k*]*S*_3_[*k*] … *S*_*n*_[*k*].

Here we are interested in estimating the probability of missassembly for each edge in the de Bruijn graph. Let *S* be a sequence (a *k*-mer) which occurs as a substring in *R a* times, i.e. the abundance of *S* in *R* is *a*. Let us assume that also the sequence *Sc* (a *k* + 1-mer), where *c* is a nucleotide, occurs as a substring in *R*. Now we are interested in whether *Sc* is a unique extension of *S* in *G* or not. If *Sc* is not a unique extension of *S* in *G*, we should not utilise the edge *Sc* in a de Bruijn graph assembly as the correctness of the edge depends on the context where *Sc* occurs and information about this context is not available in a de Bruijn graph. Note that *Sc* can be a unique extension even when *S* is a repeat in *G*. A unique extension then means that each repeat copy of *S* in *G* is followed by *c* or in other words the multiplicities of *S* and *Sc* are the same in *G*.

We are thus interested in estimating the probability that if a *k*-mer *S* is a repeat, what is the probability that an extension of *Sc* occurs with abundance at least 𝓁 in the read set when *Sc* is not a unique extension. We thus solve the following problem below:

#### Problem 1

*If S is a repeat in G, what is the probability that the abundance of Sc is larger than* 𝓁 *in R when S has at least two extensions in G, Sc and Sc*^*′*^.

This probability estimate directly gives us an estimate for an edge *Sc* to cause a misassembly.

Below we will split our analysis based on the number of repeat copies of *S* in *G*. We denote the number of repeat copies by *α* and call a repeat with *α* occurrences in *G* an *α*-repeat. Previous work exists to compute estimates for the proportion of *k*-mers that are *α*-repeats based on the abundance histogram of *k*-mers. We will utilise the Detox [13] method to compute these estimates.

### 2.2 Probability of misassembly for repeated sequences

We assume here that all sequences that occur with frequency at least 𝓁_*e*_ in the read set *R* are truly subsequences of the genome *G*. However, some of the subsequences of the genome *G* might not occur in *R*. We now want to analyse what is the probability of *Sc* not being a unique extension of *S* in *G* if the frequency of *S* and *Sc* are above some threshold 𝓁 ≥ 𝓁_*e*_ in *R*.

If the sequence *S* is not a repeat in *G*, then the probability that *Sc* is not a unique extension of *S* in *G* is 0 as there is a single occurrence of *S* in *G*. Next we will analyse the probability of an extension *Sc* of *S* not to be unique in *G* when *S* is a repeat and the frequency of *S* and *Sc* in *R* is above some threshold 𝓁 ≥ 𝓁_*e*_.

We start with the simplest scenario where *S* has exactly two occurrences in *G* and then generalise this to any *α*-repeat.

#### 2.2.1 2-repeats

The analysis given here is a slight adaptation of the analysis given in [2].

Let us assume that *S* is a repeat that occurs two times in the genome *G*. Thus there are potentially two extensions of *S* in *G*. Let us denote these by *Sc*_1_ and *Sc*_2_. Furthermore, let the abundance of *S* in the read set be *a*. If the extensions are not equal (i.e. *c*_1_≠ *c*_2_), then the extension of *Sc*_*i*_ is not unique in *G*.

Let 𝓁 *> a/*2. Now the probability that the abundance of *Sc*_1_ is at least 𝓁 is

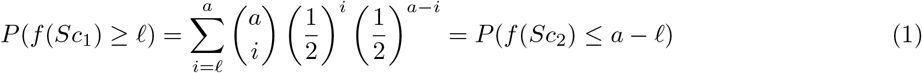

Using Chernoff’s bound^1^, we get

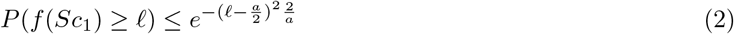

where we have utilized the fact that *f* (*Sc*_1_) ≥ 𝓁 ⇔ *f* (*Sc*_1_) ≥ (1 + *δ*)*a/*2 where *δ* = 2𝓁*/a* − 1.

Since there are two possible extensions of *S* in *G*, then the probability that an extension of *S* has an abundance above 𝓁 in *R* is

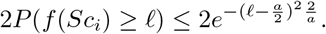

#### 2.2.2 *α*-repeats

Let us now assume that *S* is an *α*-repeat, i.e. *S* occurs in *G α* times. *S* then has *α* occurrences and *α* extensions in *G*: *Sc*_1_, *Sc*_2_, … , *Sc*_*α*_. In the worst case *α* − 1 of these extensions are equal and one is different, i.e. *c*_1_ ≠ *c*_2_ = *c*_3_ = … = *c*_*α*_. Since *S* is an *α*-repeat, we require that its extensions occur 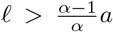 times in the read set, where *a* is the abundance of *S* in the read set. For a lower threshold we would be very likely to ignore at least one extension in *G*. Now the probability that the abundance of *Sc*, where *c* = *c*_2_ = *c*_3_ = … = *c*_*α*_, is at least 𝓁 is

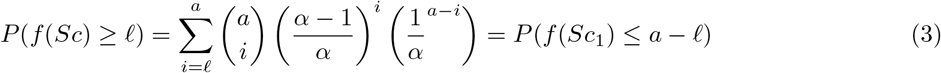

Using Chernoff’s bound^2^:

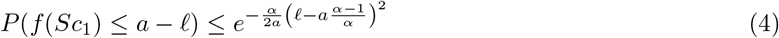

where we have utilized the fact that 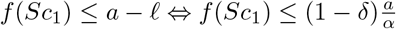 , where 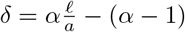.

Since there are *α* possible extensions of *S* in *G* for which the above scenario could hold, then the probability that an extension of *S* has an abundance above 𝓁 in *R* is

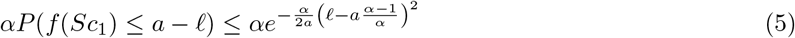

#### 2.2.3 Putting it all together

Now that we have an estimate for the proportions of *α*-repeats for different *α* and the estimates for an extension of *S* not to be unique in *G* given that *S* is an *α*-repeat, we can compute the total probability that an extension of *S* is not unique:

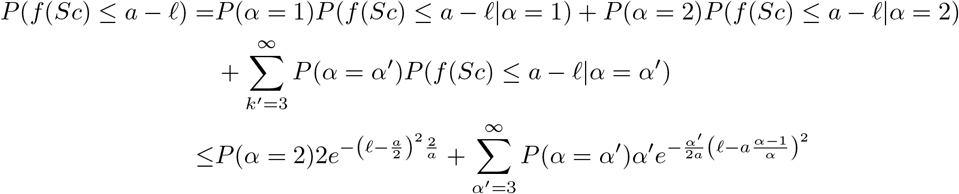

### 2.3 Integrating the analysis in de Bruijn graph based assembler

We will now use the results of the above analysis to create a de Bruijn graph based assembly where the probability of misassembly at each position is bounded by a constant *ε*. We call the resulting tool Sequence Assembly avoiding MisAssemblies (SAMA).

First we construct the de Bruijn graph using BCALM 2 [1]. We then use Detox [13] to estimate for each abundance level *a* and each *α* the probability that a *k*-mer appearing *a* times in the reads repeats *α* times in the genome. In practice, Detox limits the values of *α* in the range [0, 5]. Now we have all that is needed for the analysis presented above.

Given a *k*-mer *S* that is a node of the de Bruijn graph, each potential edge originating at that node presents an extension of *S* by one character. The above analysis gives us the probability that such an extension is not a unique extension of *S* in the genome given the abundance *a* of *S* and the abundance *a*_*e*_ of *Sc*. In particular we can now compute a lower bound for the abundance of the extension given the abundance of a *k*-mer and these can then be applied to any *k*-mer with the given abundance. Hence the next step is to compute such thresholds for each abundance of a *k*-mer. We thus iterate over all abundances of *k*-mers and for each abundance we find the lowest abundance of an extension for which the probability of an extension not being unique is less than the parameter *ε*, our probability threshold for misassembly.

The final step is to generate contigs where each de Bruijn graph edge included in the contigs meets the abundance threshold. We note that an edge between two de Bruijn graph nodes *S* and *S*^*′*^ can meet the abundance threshold in one direction but not in the other. This commonly happens at the boundary of repeats, where e.g. *S* is not a repeat *k*-mer but *S*^*′*^ is, and at the boundary of sequencing errors, where e.g. *S* is an erroneous *k*-mer but *S*^*′*^ is not. Then we can extend *S* towards *S*^*′*^ uniquely but we cannot extend *S*^*′*^ uniquely towards *S*. When generating contigs we only utilize edges which are unique extensions in both directions. We further note that the thresholds computed in the previous stage are always larger than half of the abundance of the node and therefore, at most one edge out of the node can meet the abundance threshold. Therefore, if using only such edges there is no branches to resolve.

We use the following procedure to traverse the paths of the de Bruijn graph to generate contigs. We ignore nodes with an abundance lower than five as these are likely to be results of sequencing errors. We start the traversal in some arbitrary node. We first traverse edges to the right from the chosen node as long as we find edges that meet the abundance threshold. Then we do a similar traversal to the left. During the traversal we mark each visited node to avoid producing duplicate contigs. When both traversals are complete, we produce a contig corresponding to the traversed path. We then iterate and start a new traversal from an arbitrary node that has not yet been marked (i.e. is not part of a previously generated contig). The contig generation finishes when all nodes have been marked.

## 3 Experiments

### 3.1 Datasets and methods

We ran experiments on real HiFi reads from *E. coli* and real Illumina reads from *S. aureus*. Both datasets were downsampled to the desired coverage levels. To investigate the performance of our model and other assemblers on different coverage levels, we produced three datasets of the *E. coli* dataset Ecoli20x, Ecoli40x, and Ecoli80x with coverages 20x, 40x, and 80x, respectively. The *S. aureus* was downsampled to 80x coverage.

Additionally, we simulated two datasets on human chromosome 21 with Art [3]: HumanChr21-40x with 40x coverage and HumanChr21-60x with 60x coverage. The details of the datasets are in Table 1.

**Table 1:**
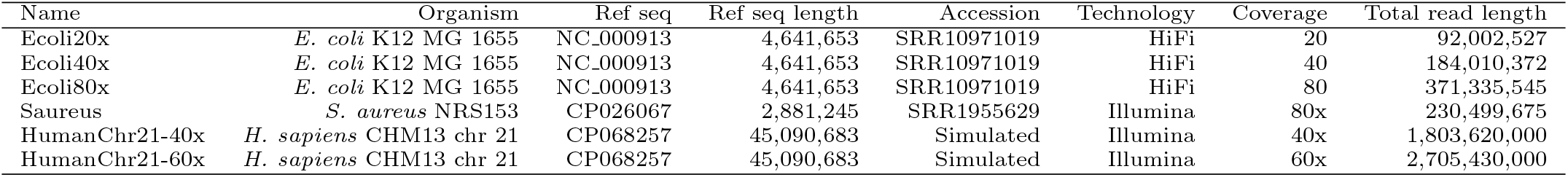
Details of the datasets. The *E. coli* and *S. aureus* datasets have been downsampled to achieve the desired coverage. The human chromosome 21 datasets are simulated with ART [3].

We compared SAMA to unitigs produced by BCALM 2 [1] and contigs produced by SPAdes [9] without error correction and before repeat resolution (which reuses the reads by mapping them on the de Bruijn graph using then the full read information to resolve repeats). BCALM 2 thus represents traditional unitigs and SPAdes contigs results of an assembly algorithm applying tip and bulge removal.

We ran all methods with two values of *k*, 31 and 63. SAMA was run with six values for the misassembly threshold *ε*: 0.4, 0.2, 0.1, 0.01, 0.001, and 0.0001. BCALM 2 was ran with three abundance thresholds: 5, 10, and 15. SPAdes was ran with the `--only-assembler` option to disable error correction and the contigs saved in the `before rr.fasta` file were analysed. We used QUAST [6] to produce statistics on the produced assemblies. All experiments were run on a computer with 32 GB of memory and 4 CPUs each with 2 cores.

### 3.2 Abundance thresholds

We first examined the abundance thresholds produced by our analysis. The thresholds as a function of *k*-mer abundance for the *E. coli, S. aureus*, and human chromosome 21 datasets are shown in Figures 1, 2, and 3, respectively. These thresholds thus indicate what is the minimum abundance of a *k* + 1-mer for it to be considered as a unique extension of a *k*-mer.

**Figure 1:**
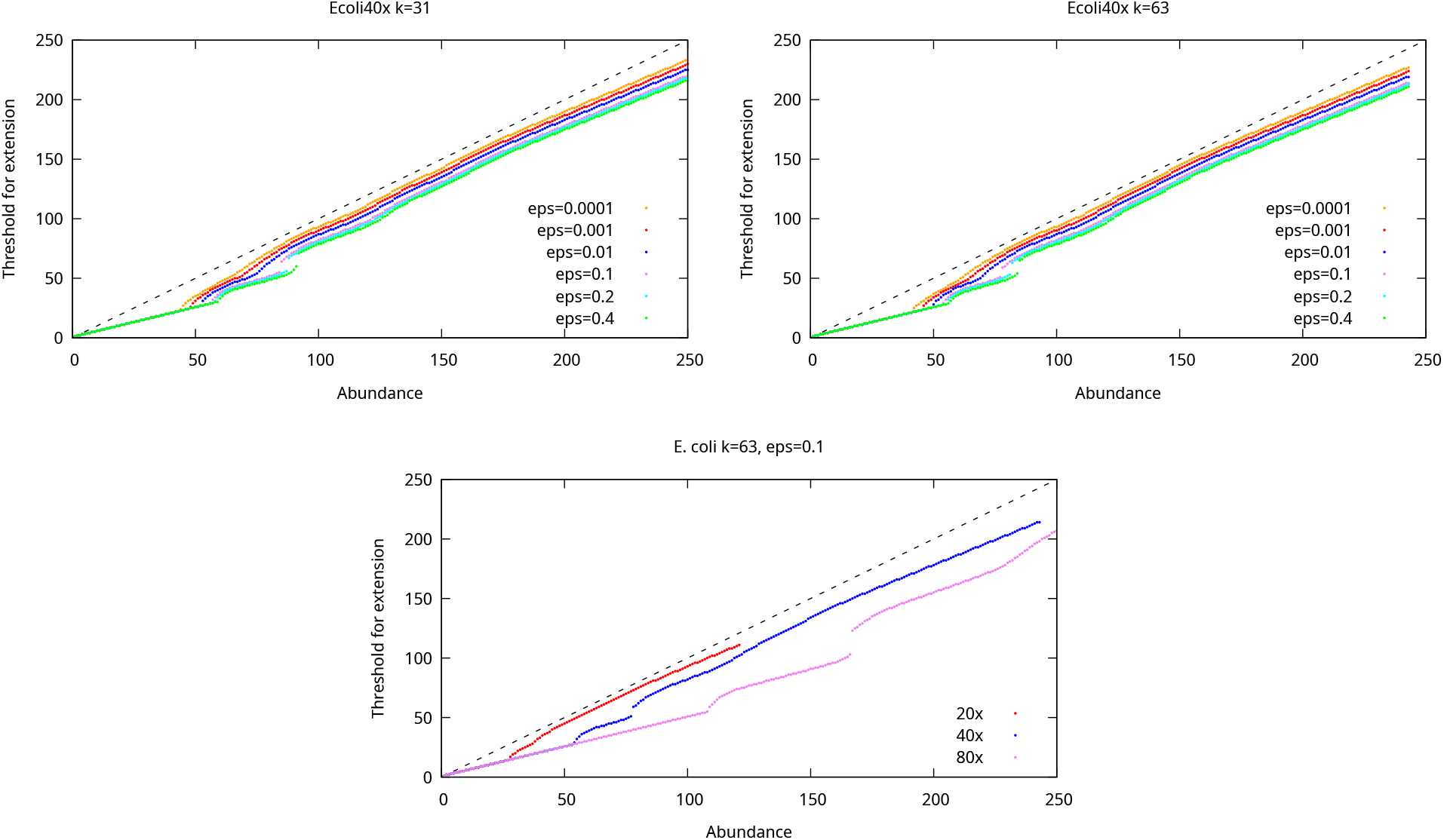
The threshold for extension as a function of *k*-mer abundance when the allowed probability for misassembly is varied (top) and when the coverage of the dataset is varied (bottom) in the *E. coli* datasets. The dashed line shows the maximum possible threshold. Missing dots indicate *k*-mer abundances where no threshold guaranteeing the desired misassembly probability can be found.

**Figure 2:**
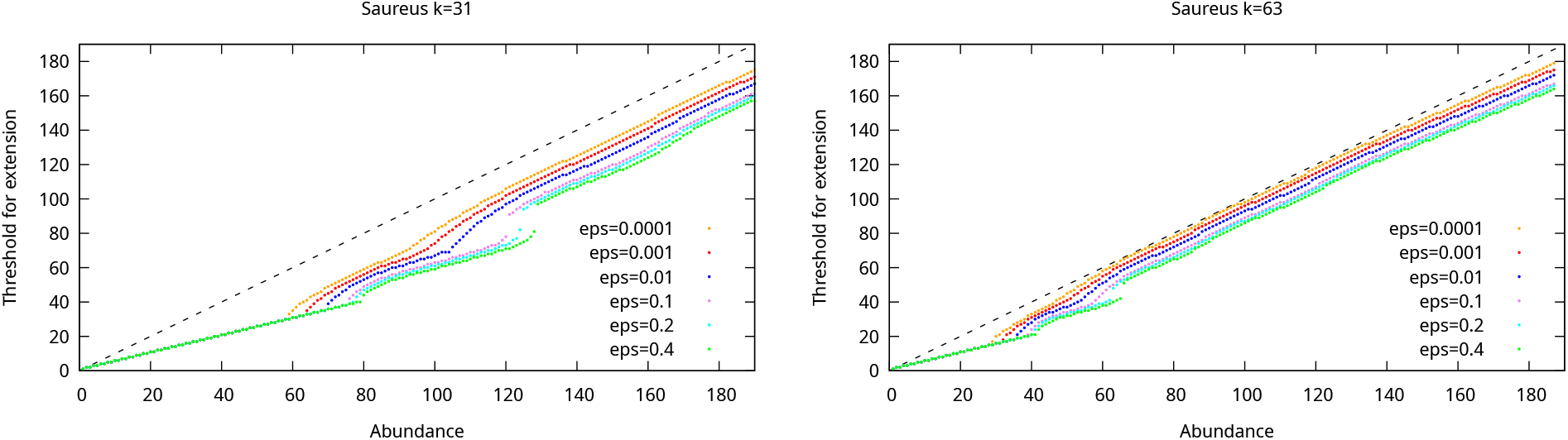
The threshold for extension as a function of *k*-mer abundance when the allowed probability for misassembly is varied for the Saureus dataset. The dashed line shows the maximum possible threshold. Missing dots indicate *k*-mer abundances where no threshold guaranteeing the desired misassembly probability can be found.

**Figure 3:**
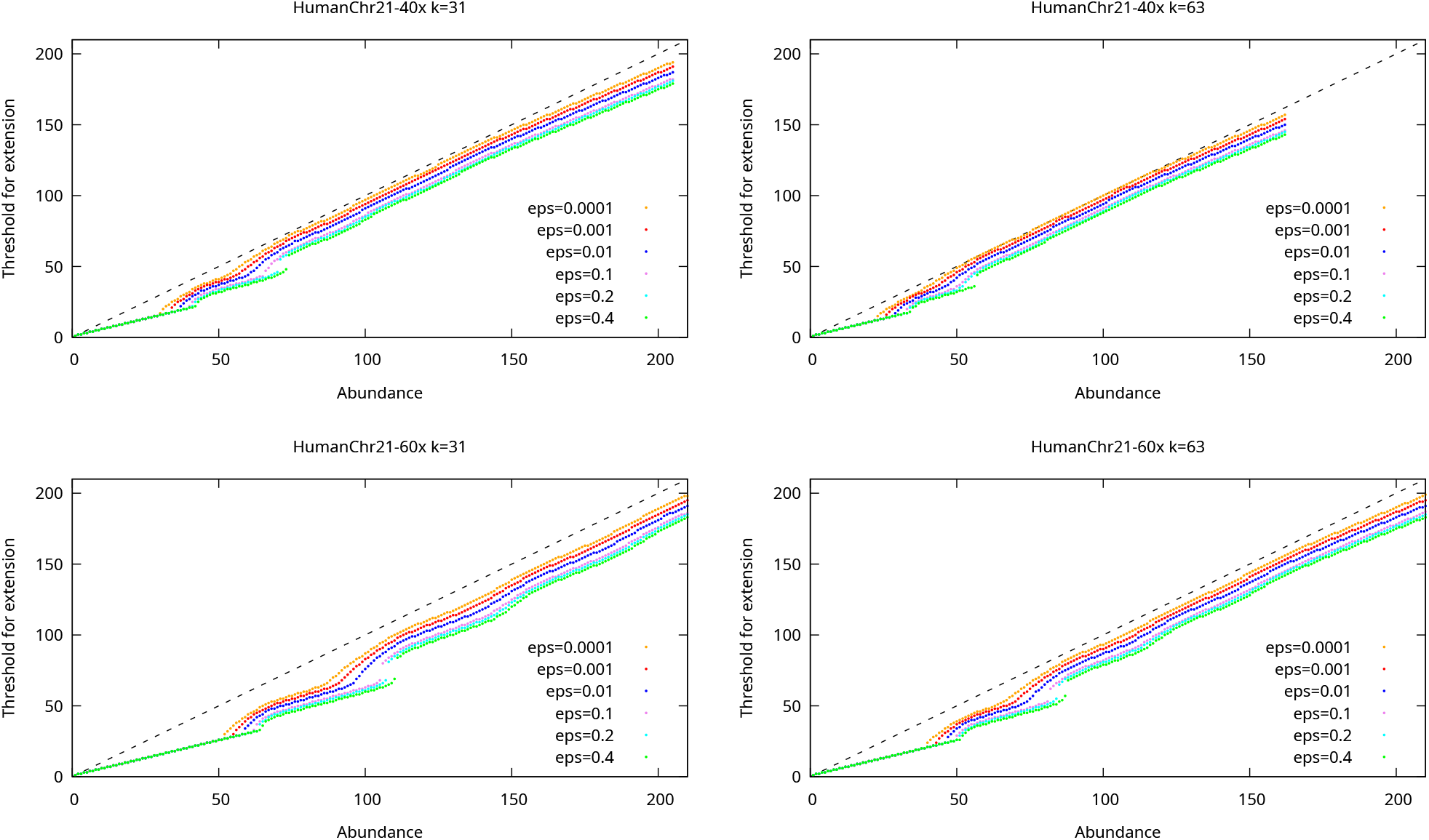
The threshold for extension as a function of *k*-mer abundance when the allowed probability for misassembly is varied for the HumanChr21-40x dataset (top) and Human Chr21-60x dataset (bottom). The dashed line shows the maximum possible threshold. Missing dots indicate *k*-mer abundances where no threshold guaranteeing the desired misassembly probability can be found.

For low abundance *k*-mers the thresholds are close to half the abundance of the *k*-mers because these *k*-mers are likely to be unique in the genome and for the unique *k*-mers probability of misassembly in our model is 0. For the higher abundance *k*-mers, the thresholds are close to the abundance of the *k*-mers due to most of these *k*-mers originating from repeats and then a lot of evidence is needed to determine that the extension is likely to be unique. However, the increase in the threshold is not even which is due to the proportions of different *α*-repeats changing when the abundance of *k*-mers increases. We also see that the thresholds increase as *ε* increases. When the probability of misassembly is set low, we find some *k*-mer abundances for which no abundancy threshold exist such that we could guarantee a low enough misassembly probability.

### 3.3 Comparison to other methods

We computed the following statistics on all the assemblies produced by SAMA, BCALM 2, and SPAdes: **#contigs** : The number of contigs longer than 100 bp in the assembly.

#### NGA50

The contigs are first aligned on the reference genome. The alignments are then ordered from largest to smallest. The NGA50 metric is the length of the alignment where the cumulative length of the alignments exceeds half the genome length.

#### Misassemblies

The number of misassemblies as reported by QUAST.

#### Genome fraction

The percentage of nucleotides in the genome that are covered by at least one contig. Table 2 shows the statistics of the assemblies produced by the various methods on the *E. coli* datasets.

**Table 2:** Assembly statistics for the *E. coli* datasets.

| Dataset | Method | Parameters | #contigs ($\geq 100$ bp) | NGA50 | Misassemblies | Genome fraction (%) |
| --- | --- | --- | --- | --- | --- | --- |
| Ecoli20x | SAMA | k=31 $\varepsilon=0.4$ | 343 | 21,181 | 0 | 97.0 |
| | SAMA | k=31 $\varepsilon=0.2$ | 349 | 21,081 | 0 | 97.0 |
| | SAMA | k=31 $\varepsilon=0.1$ | 351 | 21,081 | 0 | 97.0 |
| | SAMA | k=31 $\varepsilon=0.01$ | 358 | 20,068 | 0 | 96.9 |
| | SAMA | k=31 $\varepsilon=0.001$ | 379 | 18,858 | 0 | 96.8 |
| | SAMA | k=31 $\varepsilon=0.0001$ | 498 | 15,051 | 0 | 96.4 |
| | SAMA | k=63 $\varepsilon=0.4$ | 131 | 65,330 | 0 | 97.6 |
| | SAMA | k=63 $\varepsilon=0.2$ | 131 | 65,330 | 0 | 97.6 |
| | SAMA | k=63 $\varepsilon=0.1$ | 135 | 60,614 | 0 | 97.6 |
| | SAMA | k=63 $\varepsilon=0.01$ | 138 | 59,680 | 0 | 97.5 |
| | SAMA | k=63 $\varepsilon=0.001$ | 157 | 51,643 | 0 | 97.5 |
| | SAMA | k=63 $\varepsilon=0.0001$ | 313 | 34,150 | 0 | 96.6 |
|  | BCALM2 | k=31 a=5 | 747 | 13,532 | 0 | 97.6 |
|  | BCALM2 | k=31 a=10 | 1,101 | 15,302 | 0 | 96.3 |
|  | BCALM2 | k=31 a=15 | 4,188 | 2,413 | 0 | 78.4 |
|  | BCALM2 | k=63 a=5 | 874 | 31,596 | 0 | 98.7 |
|  | BCALM2 | k=63 a=10 | 1,660 | 26,917 | 0 | 97.1 |
|  | BCALM2 | k=63 a=15 | 6,231 | 950 | 0 | 77.2 |
|  | SPAdes | k=31 | 469 | 22,175 | 0 | 97.1 |
|  | SPAdes | k=63 | 259 | 78,620 | 0 | 98.2 |
| Ecoli40x | SAMA | k=31 $\varepsilon=0.4$ | 334 | 22,902 | 1 | 97.1 |
| | SAMA | k=31 $\varepsilon=0.2$ | 339 | 22,902 | 1 | 97.1 |
| | SAMA | k=31 $\varepsilon=0.1$ | 347 | 22,580 | 0 | 97.1 |
| | SAMA | k=31 $\varepsilon=0.01$ | 348 | 21,793 | 0 | 97.0 |
| | SAMA | k=31 $\varepsilon=0.001$ | 350 | 21,793 | 0 | 97.0 |
| | SAMA | k=31 $\varepsilon=0.0001$ | 351 | 21,541 | 0 | 96.9 |
| | SAMA | k=63 $\varepsilon=0.4$ | 124 | 73,700 | 0 | 97.7 |
| | SAMA | k=63 $\varepsilon=0.2$ | 126 | 73,700 | 0 | 97.7 |
| | SAMA | k=63 $\varepsilon=0.1$ | 130 | 73,700 | 0 | 97.7 |
| | SAMA | k=63 $\varepsilon=0.01$ | 128 | 73,700 | 0 | 97.6 |
| | SAMA | k=63 $\varepsilon=0.001$ | 133 | 59,680 | 0 | 97.6 |
| | SAMA | k=63 $\varepsilon=0.0001$ | 141 | 57,934 | 0 | 97.6 |
|  | BCALM2 | k=31 a=5 | 2112 | 4,082 | 0 | 97.5 |
|  | BCALM2 | k=31 a=10 | 576 | 18,227 | 0 | 97.6 |
|  | BCALM2 | k=31 a=15 | 541 | 21,235 | 0 | 97.7 |
|  | BCALM2 | k=63 a=5 | 3751 | 5,702 | 0 | 98.9 |
|  | BCALM2 | k=63 a=10 | 583 | 59,552 | 0 | 98.7 |
|  | BCALM2 | k=63 a=15 | 542 | 63,602 | 0 | 98.7 |
|  | SPAdes | k=31 | 485 | 22,175 | 0 | 97.8 |
|  | SPAdes | k=63 | 274 | 78,620 | 0 | 98.4 |
| Ecoli80x | SAMA | k=31 $\varepsilon=0.4$ | 342 | 23,326 | 0 | 97.0 |
| | SAMA | k=31 $\varepsilon=0.2$ | 344 | 22,902 | 0 | 97.0 |
| | SAMA | k=31 $\varepsilon=0.1$ | 344 | 22,902 | 0 | 97.0 |
| | SAMA | k=31 $\varepsilon=0.01$ | 344 | 22,902 | 0 | 97.0 |
| | SAMA | k=31 $\varepsilon=0.001$ | 345 | 22,902 | 0 | 97.0 |
| | SAMA | k=31 $\varepsilon=0.0001$ | 346 | 22,173 | 0 | 97.0 |
| | SAMA | k=63 $\varepsilon=0.4$ | 129 | 78,618 | 0 | 97.7 |
| | SAMA | k=63 $\varepsilon=0.2$ | 128 | 78,618 | 0 | 97.7 |
| | SAMA | k=63 $\varepsilon=0.1$ | 127 | 78,618 | 0 | 97.7 |
| | SAMA | k=63 $\varepsilon=0.01$ | 126 | 78,618 | 0 | 97.7 |
| | SAMA | k=63 $\varepsilon=0.001$ | 126 | 78,618 | 0 | 97.7 |
| | SAMA | k=63 $\varepsilon=0.0001$ | 126 | 78,618 | 0 | 97.6 |
|  | BCALM2 | k=31 a=5 | 8,019 | 813 | 0 | 95.4 |
|  | BCALM2 | k=31 a=10 | 1,465 | 6,189 | 0 | 97.5 |
|  | BCALM2 | k=31 a=15 | 653 | 16,190 | 0 | 97.6 |
|  | BCALM2 | k=63 a=5 | 19,389 | 986 | 0 | 99.0 |
|  | BCALM2 | k=63 a=10 | 2,275 | 9,466 | 0 | 98.8 |
|  | BCALM2 | k=63 a=15 | 723 | 41,980 | 0 | 98.8 |
|  | SPAdes | k=31 | 485 | 22,175 | 0 | 97.8 |
|  | SPAdes | k=63 | 268 | 78,620 | 0 | 98.3 |

Similarly, Table 3 shows the statistics of the assemblies produced by the methods on the Saureus and human chromosome 21 datasets.

**Table 3:**
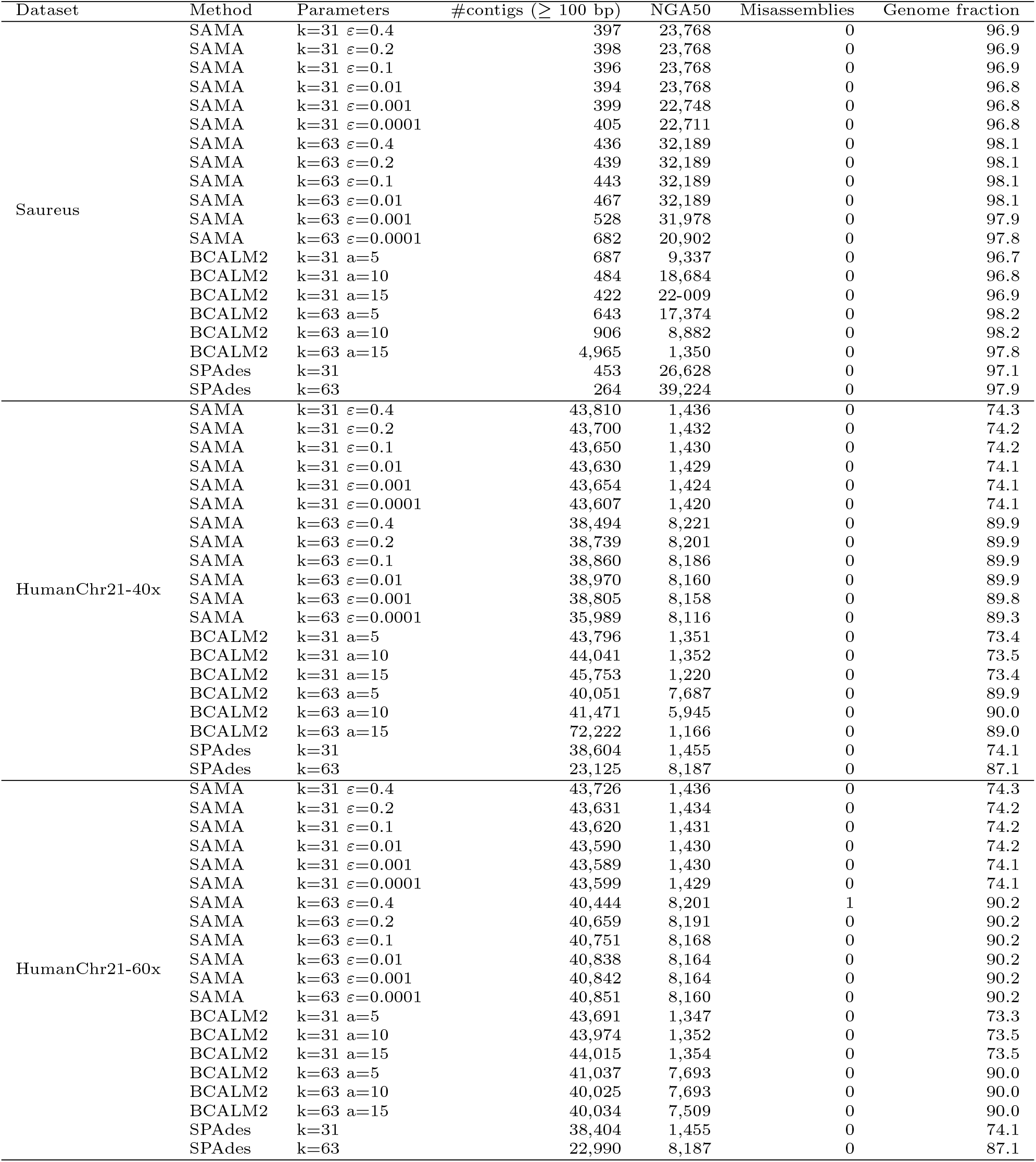
Assembly statistics for the Saureus and HumanChr21 datasets.

We see that SAMA produces more contiguous assemblies, i.e. lower number of contigs, higher NGA50, and higher genome fraction, when we allow a higher probability for misassemblies. SAMA produces misassemblies only on the Ecoli40x and HumanChr21-60x datasets when the allowed probability for misassemblies is high. When the allowed probability of misassemblies is less than or equal to 0.1 there are no misassemblies on any of the datasets. We also see that when we attempt to produce assemblies with very small probability of misassembly on the low coverage Ecoli20x dataset, SAMA produces fragmented assemblies as the dataset does not provide enough data for such accurate assemblies.

SAMA produces more contiguous assemblies than the unitigs produced by BCALM 2 when the allowed probability for misassemblies is set appropriately. When the coverage of the dataset is high enough for accurate enough probability estimates, SAMA produces assemblies with similar contiguity characteristics as SPAdes and with no misassemblies. When coverage is lower (Ecoli20x, Ecoli40x, Saureus, HumanChr21-40x datasets), SPAdes produces somewhat more contiguous assemblies. On the other hand, on the more complex human chromosome 21 datasets, SAMA produces more contiguous assemblies than SPAdes. It is worth noting that although the Saureus dataset has nucleotide coverage of 80x, the read length is only 101 bp, and so the coverage of *k*-mers is only 56x and 31x when *k* is 31 and 63, respectively.

Table 4 shows the runtime and peak memory usage of the tools. The statistics for SAMA include running BCALM 2 and Detox as they part of the needed pipeline. The comparison against SPAdes is not entirely fair as it was not possible to disable running the repeat resolution code which uses a significant amount of resources. We see that SAMA is competitive in both runtime and memory usage.

**Table 4:** Runtime and memory usage.

| Dataset | Method | Parameters | Runtime (min) | Memory Usage (GB) |
| --- | --- | --- | --- | --- |
| Ecoli20x | SAMA | k=31 $\epsilon=1e-01$ | 0.90 | 1.27 |
| | SAMA | k=63 $\epsilon=1e-01$ | 0.87 | 1.34 |
|  | BCALM2 | k=31 a=10 | 0.23 | 0.58 |
|  | BCALM2 | k=63 a=5 | 0.32 | 0.87 |
|  | SPAdes | k=31 | 0.50 | 0.79 |
|  | SPAdes | k=63 | 0.84 | 1.53 |
| Ecoli40x | SAMA | k=31 $\epsilon=1e-01$ | 1.17 | 1.45 |
| | SAMA | k=63 $\epsilon=1e-01$ | 1.34 | 1.66 |
|  | BCALM2 | k=31 a=15 | 0.35 | 0.98 |
|  | BCALM2 | k=63 a=15 | 0.44 | 1.55 |
|  | SPAdes | k=31 | 0.97 | 1.54 |
|  | SPAdes | k=63 | 1.64 | 2.21 |
| Ecoli80x | SAMA | k=31 $\epsilon=1e-01$ | 1.88 | 1.94 |
| | SAMA | k=63 $\epsilon=1e-01$ | 2.22 | 2.88 |
|  | BCALM2 | k=31 a=15 | 0.54 | 1.71 |
|  | BCALM2 | k=63 a=15 | 0.68 | 2.88 |
|  | SPAdes | k=31 | 1.83 | 2.21 |
|  | SPAdes | k=63 | 3.29 | 2.22 |
| Saureus | SAMA | k=31 $\epsilon=1e-01$ | 0.64 | 0.88 |
| | SAMA | k=63 $\epsilon=1e-01$ | 0.45 | 0.81 |
|  | BCALM2 | k=31 a=15 | 0.28 | 0.76 |
|  | BCALM2 | k=63 a=5 | 0.28 | 0.76 |
|  | SPAdes | k=31 | 0.90 | 1.34 |
|  | SPAdes | k=63 | 0.66 | 1.45 |
| HumanChr21-40x | SAMA | k=31 $\epsilon=1e-01$ | 23.64 | 11.40 |
| | SAMA | k=63 $\epsilon=1e-01$ | 14.14 | 11.69 |
|  | BCALM2 | k=31 a=10 | 2.86 | 3.23 |
|  | BCALM2 | k=63 a=5 | 3.32 | 3.34 |
|  | SPAdes | k=31 | 132.93 | 18.79 |
|  | SPAdes | k=63 | 32.77 | 6.92 |
| HumanChr21-60x | SAMA | k=31 $\epsilon=1e-01$ | 26.77 | 12.69 |
| | SAMA | k=63 $\epsilon=1e-01$ | 19.52 | 12.81 |
|  | BCALM2 | k=31 a=15 | 3.78 | 4.17 |
|  | BCALM2 | k=63 a=5 | 4.28 | 4.30 |
|  | SPAdes | k=31 | 167.78 | 25.62 |
|  | SPAdes | k=63 | 47.21 | 9.53 |

## 4 Conclusions and further work

We have presented a model to estimate the probability of misassembly at each position of a de Bruijn graph based assembly. As far as we know, this is the first model that can assign such a probability to each position in the assembly also taking into account missing data. We applied our model to produce contigs with correctness guarantee and showed that this leads to a practical method. Here we applied the model to produce an assembly where the probability of misassembly at each position is bounded. However, this information has further potential applications in the downstream analysis of a genome and in any analysis method working directly on the de Bruijn graph. Our current model is limited to single haploid genomes. It would be interesting to extend the ideas presented here to diploid genomes as well as viral and bacterial metagenomics.

## Footnotes

1 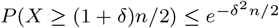 for *X* that is distributed as Bernoulli(1/2)

2 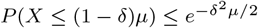 for *X* that is Binomial(*n, p*)

